# Feature Selection Using Lasso Regression Enhances Deep Learning Model Performance For Diagnosis Of Lung Cancer from Transcriptomic Data

**DOI:** 10.1101/2024.05.01.592076

**Authors:** Souvik Guha

## Abstract

Cancer is a genetic disease where gene mutations are pivotal in disease initiation and pathophysiology. The gene expression profile follows a specific pattern exclusive to each cancer which can be utilized for early and accurate diagnosis. Microarray techniques have emerged as powerful tools capable of simultaneously capturing the expression profiles of thousands of genes. However, because of the high dimensionality of the produced transcriptome data, analysis of the resulting datasets is challenging. Recent advancements in Artificial Intelligence (AI) techniques like Machine Learning (ML) and Deep Learning can be instrumental in efficiently processing these high-dimensional datasets. LASSO-regression is a ML technique that can help to rank the features which could help in feature selection leading to dimensionality reduction. Deep Learning is one of the most sophisticated ML techniques that can process high-dimensional data owing to the presence of more number of hidden layers in its neural network. We designed a Deep Neural Network (DNN) classifier model fused with a LASSO-based significant feature extractor for classifying the gene expression dataset containing a total of 51 samples of which 24 samples are of lung cancer patients and the remaining 27 samples are of normal individuals. A LASSO regression model was implemented to identify the genes that played a significant role in the classification. These significant gene expressions were then fed into a convergent Deep Neural Architecture. The classifier was trained with 70% data and the rest 30% was used for validation. The proposed classifier proved to provide better classification as compared to LASSO regression and DNN used individually. The two classes were classified with an average accuracy of 96.25%, average precision of 99.67%, average specificity of 99.45% and average sensitivity of 91.73% measured over thirty independent assessments. In some cases, the model was able to obtain a classification accuracy of 100%. This could open the path to early and better diagnosis of cancers from transcriptome data.

## 1. Introduction

An individual’s genetic information is stored in their genes. Any medical condition resulting from abnormalities in the expression of a person’s genes is classified as a genetic disease. It can result from chromosome damage, several gene mutations, a single gene mutation, a mix of gene mutations, and many environmental conditions. Cancer is also often caused as a result of certain genetic diseases or abnormalities that can pass from parent to child. Globally, lung cancer is the most prevalent cancer among those diagnosed. With 1.38 million cancer deaths annually globally, it is also the leading cause of cancer-related mortality [1]. With an anticipated 1.8 million fatalities (18%), lung cancer continued to be the most common cause of cancer-related mortality. Colorectal (9.4%), liver (8.3%), stomach (7.7%), and female breast cancers (6.9%) were the next most common causes [2]. Several factors can play a role in the development of lung cancer. The main risk factor for lung cancer is tobacco use, and smoking cigarettes is directly responsible for a significant percentage of all cases of lung cancer [3]. According to World Health Organization predictions, there will be an ongoing spike in lung cancer fatalities globally, primarily due to rising tobacco use worldwide, especially in Asia. Metastasis from other tumours located in distinct parts can also initiate lung cancer. Study shows that approximately 60% of metastatic breast cancer patients suffer lung or bone metastasis in their life [4]. Another significant factor is genetics. A risk prediction model developed by *Spitz, et al* included several factors, like past smoking history, exposure to secondhand smoke, occupational exposure to asbestos and dust, and a family history of cancer. They demonstrated how a family history of cancer affected the risk of lung cancer in those who had never smoked, had smoked in the past, and currently smoke. The model showed a significant risk of lung cancer development in individuals who never smoked but had a family history of lung cancer [5]. Cassidy, *et al* also drew emphasis on the significantly elevated risk of lung cancer, particularly in those with a family history of early-onset (< 60 years old) lung cancer [6]. The DNA which is the inherited element is transcribed to mRNA which in turn forms the proteins. Hence the alterations in the gene expressions can be tracked by monitoring the transcriptome profile. DNA microarray technique is one of the ways of studying the expression profile of a large number of genes simultaneously. It is a highly efficient technique to detect any mutations present in the DNA of an individual. Microarray techniques have been widely being used for diagnosis, prognosis and development of better therapeutics for several types of cancers [7]. This technique hence could aid in the development of novel automated diagnostic techniques that can help in the early diagnosis of several types of cancer from the transcriptomic profile.

The output data of the microarray analysis are high dimensional matrices that contain expression profiles of thousands of genes belonging to a large number of samples. Efficient processing and feature extraction hence remain a challenge. The recent advancement in the field of computational techniques and the emergence of Artificial Intelligence driven Machine Learning could be instrumental in handling the processing of these high dimensional data. Machine learning techniques like Lasso regression and advanced techniques like Deep Learning are widely being used for the processing of biological data [8]. Within machine learning-based biomedical data analysis, gene expression data research is one of the most prominent fields. Higher dimensional data could result in poor performance in ML algorithms. Higher dimensionality can lead to overly complex models that fit the noise instead of the underlying pattern. The model’s capacity to generalize to new data is weakened as a result. In this work, we have demonstrated that dimensional reduction by feature extraction could marginally improve the performance of the Deep Learning classifier for classifying between cancer and healthy samples. We have used the Lasso regression classifier to identify the significant genes from the input dataset. These genes were then fed into a Deep Neural Engine Classifier model that did the final classification. The model parameters were compared with feature-reduced and without feature-reduced datasets. The former shows better classification parameters. This could potentially lead to the development of better classifier models for diagnostic purposes.

## 2. Literature Review

The recent advancement in AI has increased the usage of techniques like ML and DL in the processing of biological data. The principal aim of using these techniques is to improve the performance of these classification models by improving their classification parameters and decreasing the error. Feature selection is important in the analysis of high-dimensional datasets. Sharma, *et al*. have used a null space-based feature extraction technique. The model removes the redundant genes by applying the information of null space of scatter matrices [9]. After removing the insignificant features three different classifiers such as Support Vector Machine (SVM), Nave Bayes and Linear Discriminant Analysis (LDA) were employed. Vasighizaker et al. suggested a one-class SVM classifier that was used in gene expression data to identify AML samples. The effectiveness of their suggested classifier was tested using various kernel function types, and it was found that a linear kernel produced superior accuracy. Additionally, they asserted that their conclusion was superior to other models by comparing it with a few older, conventional classification models [10]. Jin X, *et al*. addressed the high-dimension challenge by implementing the chi-square feature selection approach. They have used the SAGE data set to use machine learning techniques in their study. They concluded from the data that SVM and Naive Bayes classifiers performed better with fewer features [11]. J. Bennet, C., *et al*. suggested combining the Based Bayes Error Filter (BBF) for attribute selection with the support vector machine-recursive feature elimination (SVM-RFE) approach. The attributes were sorted using SVM-RFE by these researchers, and any redundant sorted attributes were eliminated using BBF. Then, for classification, the SVM algorithm was applied [12]. Bouazza, Sara Haddou, *et al*. worked on employing KNN and SVM classifiers to classify cancer cases. In their study, the impacts on three gene expression profile datasets (prostate, colon, and leukaemia) were investigated utilizing a variety of attribute selection strategies, including KNN and SVM classifiers and Fisher, ReliefF, SNR, and T-Statistics. Combining the SNR attribute selection method with the SVM classifier produced the best outcomes [13]. Huynh, *et al*. have employed several Deep learning techniques for the interpretation of gene expression data. Using a convolutional neural network, the pertinent gene characteristics were retrieved [14].

From the literature, it is clear that multiple studies have been performed for gene expression classification. These works have demonstrated that feature reduction can significantly improve the performance of the classifier model by reducing the noise and overfitting simultaneously decreasing the complexity and computational time. However none of them have used the Lasso regression model for feature reduction. In our proposed model we have employed a Lasso regression model as the feature extractor and subsequently used a Convergent Deep Neural Classifier for classification.

## 3. Proposed Model

The proposed Lasso Feature extraction enhanced Deep Learning lung cancer classifier model consists of three primary blocks [Figure 1]. We took the dataset from the NCBI Gene Expression Omnibus which was then fed into a preprocessing unit that prepared all the required preprocessings. After preprocessing the data was fed into a Lasso regression-based classifier which ranked the features by defining the coefficients of all the features(genes). Only the genes having a non-zero coefficient were then fed into a Convergent Deep Learning classifier which finally performed the classification. The classification parameters were compared both for the feature reducted and the dataset with no feature reduction. The Deep Learning model performed marginally well in the feature-reduced dataset.

**Figure 1.**
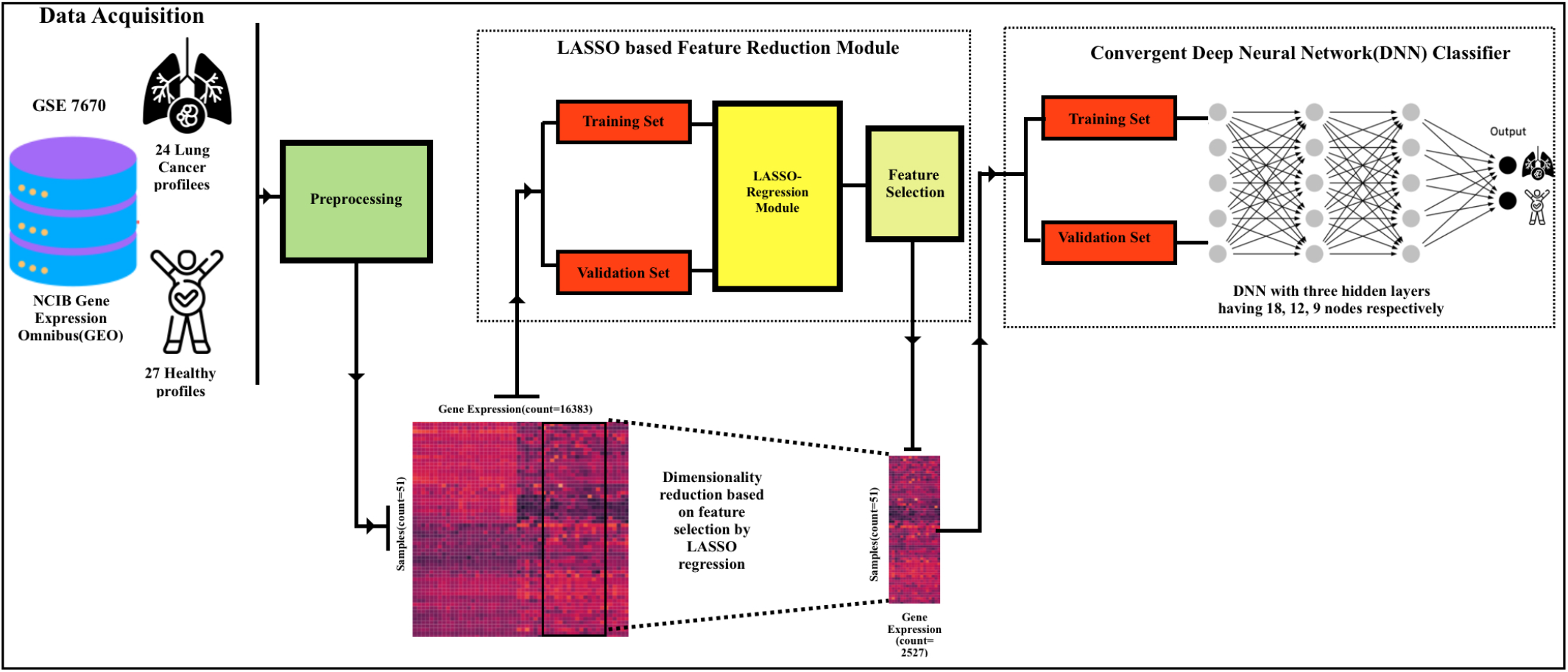
Schematics of the proposed model. The model consists of three primary modules, the Preprocessing, the Lasso based feature extraction and the Convergent Deep Neural Network Classifier.

### 3.1 Dataset Acquisition

The NCBI Gene Expression Omnibus (GEO) was queried for lung cancer datasets extensively. Finally, dataset with an accession ID GSE7670 was chosen for the study. The microarray dataset consists of a total of 16382 gene expression profiles belonging to 27 healthy individuals and 24 lung cancer patients.

### 3.2 Dataset Preprocessing

Data pre-processing operations included checking for any missing values in the dataset. The edgeR package in R was applied to the dataset to get the *log*_2_ transformation and the upper quartile (normalization) of the data. The ARSyNseq function of the NOISeq package was applied to this normalized data to perform batch correction. The probe-id of the genes was not matched with their respective gene symbols because the names of the individual genes were irrelevant to the current study, the feature extraction was performed based on the probe-id. The samples belonging to the cancer patients were labelled as “1” whereas the samples belonging to the healthy individuals were marked as “0”.

### 3.3 Dataset Preparation for Lasso Regression Classifier

The dataset is a matrix of dimensions (51×16382) representing (samples×gene expression levels). For training the model we did a 7:3 split along the samples. 70% data was used for training the model and 30% data was used for the validation purposes. The dataset was split using the train-test split function available in the sklearn package which offered shuffling of the dataset before splitting which could reduce any potential bias.[15].

### 3.4 Lasso Regression Model for Feature Selection

Lasso regression enhances the linear regression model by incorporating additional data into the model through a process called L1 regularization, which prevents overfitting. Therefore, by fitting a model with every possible predictor and using a regularization technique that lowers the coefficient estimates to zero, we may use Lasso for efficient feature extraction. In particular, the reduction objective comprises, as in the OLS regression setting, the residual sum of squares (RSS) plus the sum of the absolute values of the coefficients.

The residual sum of squares (RSS) is calculated as follows:

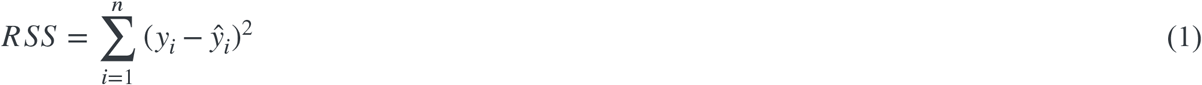

This can also be stated as:

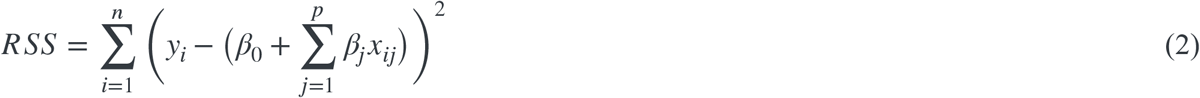

n represents the number of observations, p denotes the number of genes *x*_*ij*_ and represents the value of the jth variable for the ith observation (gene expression level), where i = 1, 2, …, n and j = 1, 2, …, p.

Within the lasso regression, the goal of minimization is transformed into:

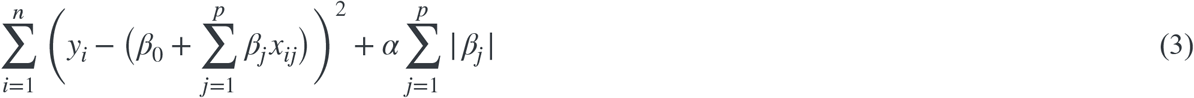

Or

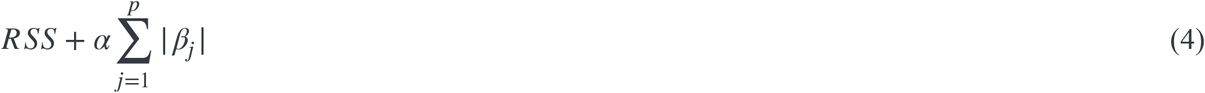

Where can take various values, depending on the dataset the value is estimated.

The advantage of Lasso regression over least squares linear regression is due to the bias-variance trade-off. Lasso can be instrumental in feature selection from a high-dimensional dataset [16]. The feature extraction was done by following steps:

**Step 1:** The optimal value was estimated using the GridSearchCV [17] module and was implemented into the Lasso module of Scikit.

**Step 2:** The model was trained with the training set.

**Step 3:** The coefficients of all the genes present in the database were calculated.

**Step 4:** The coefficients were sorted in descending order.

**Step 5:** The genes having non-zero coefficients were extracted and other gene expression profiles were removed.

**Step 6:** The model was evaluated using the validation set and its performance was evaluated, Step 1 was to be repeated if required.

### 3.5 Dataset Preparation for Deep Learning Classifier

The feature-reduced dataset that was obtained after was again split in a 7:3 ratio where 70% data was used for training the model and 30% of the data was used for validation purposes. The dataset was split using the train-test split function available in the sklearn package which offered shuffling of the dataset before splitting which could reduce any potential bias.[15].

### 3.6 Convergent Deep Learning Classifier Model

Deep Learning is one of the recently developed and one of the most advanced branches of artificial intelligence (AI) that has become a significant topic of interest due to its exceptional learning abilities within the data-driven industry for better analysis of the growing amount of generated data. Deep is a form of AI function that simulates the human brain’s ability to process data [19]. A deep neural network (DNN) implements the concept of a feed-forward neural network in which the input data is directed from the input layer to the output layer without flowing backwards [Figure 2]. At the beginning, a connection is mapped between the virtual neurons (nodes) with random values as weights. Following this, the input data is multiplied by the weights, and the result, which is between 0 and 1, is continuously evaluated during the training process. The weights are then adjusted if the model is unable to recognize the pattern [20]. The proposed DNN was implemented in Python using the Keras (https://keras.io) library package. We used three hidden layers and 140 epochs with a dropout of rate 0.2 after each layer which randomly sets input to 0 which helps to reduce the overfitting of the model. The input gene data vector of the form G = {g_1_, g_2_, g_3_, g_4_, …, g_n_} is multiplied with the weight vector of the form W = {w_1_, w_2_, w_3_, w_4_, …, w_n_}. The output O of each individual layer after multiplying the gene vector with the weight vector is given as

**Figure 2.**
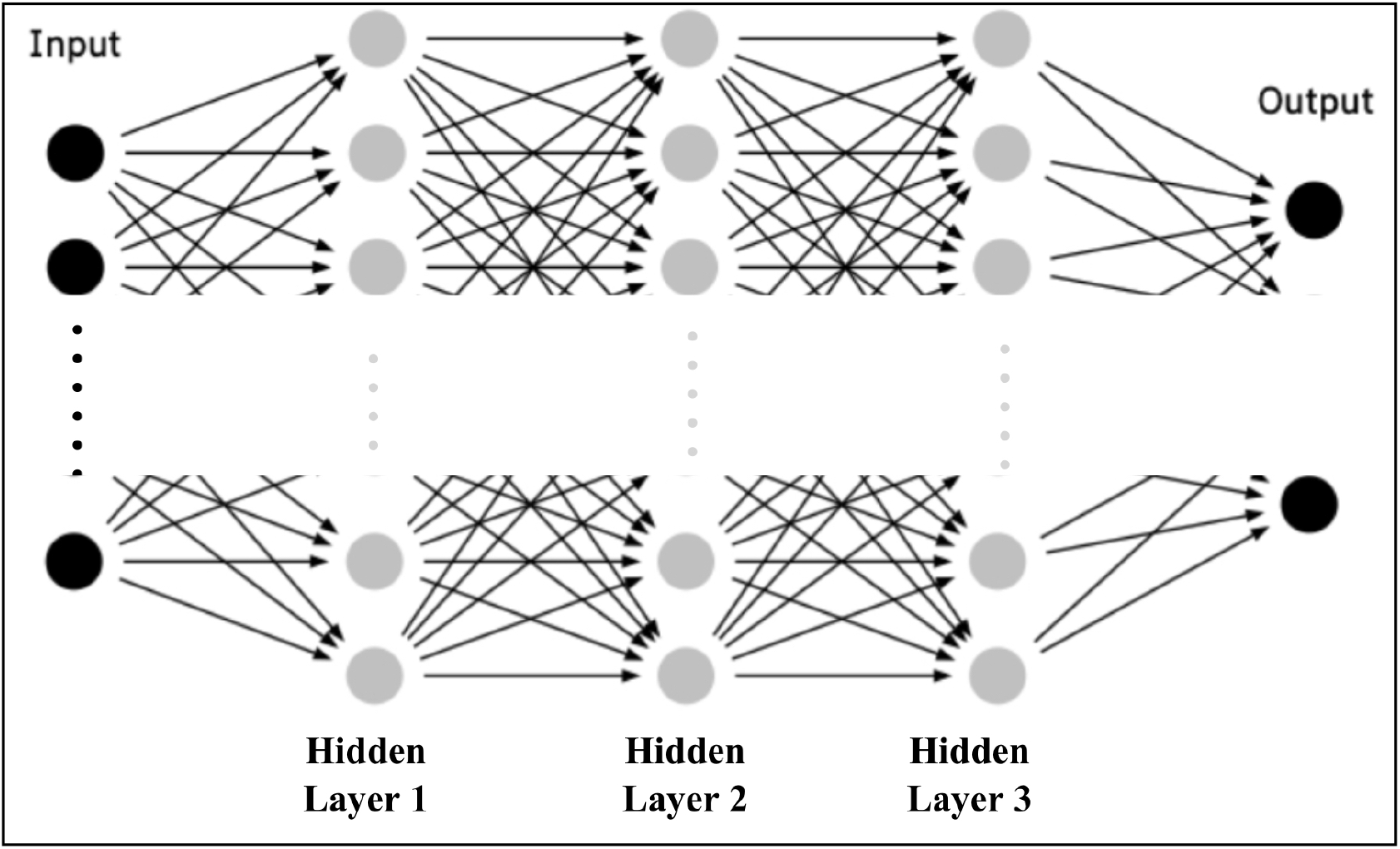
Representation of a typical DNN structure. This DNN has three hidden layers, one input layer and one output layer.

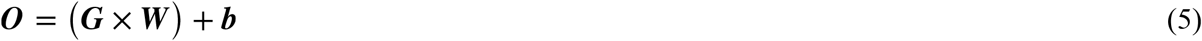

Where ***b*** is called the bias.

After multiplication of the weight vector with the input data (genes), it is passed through a nonlinear activation function which can be represented as

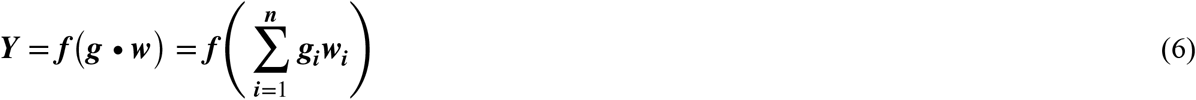

Where ***Y*** is the output of the classifier and n is the number of input features (genes). The activation function ***f*** is defined for each hidden layer and the output layer. In this model, we have used ReLu as the activation function for each hidden layer, which is defined as

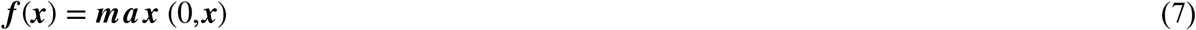

The ReLu is a very popular activation function in DNN algorithms. Training in this activation function is comparatively much faster than in other activation functions because it has the advantage of not activating all neurons simultaneously and simultaneously converting all negative inputs into 0. This ensures that the neurons do not get activated. For the output layer, we have implemented the Sigmoid activation function which is defined as

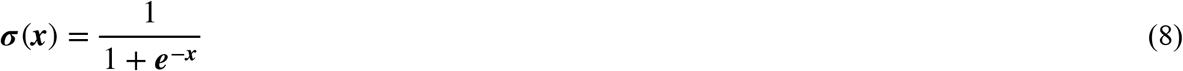

The Sigmoid activation function exists between (0,1) hence it is suitable for binary classifications., it also has the advantage of faster training like the ReLu activation function though it is only suitable for shallow neural networks.

### 3.7 Model Evaluation

The models were executed 30 times with independent training and validation sets and their performance was calculated from constructing the corresponding Confusion Matrices. The metrics were taken as the average over 30 iterations. The evaluation metric of the models is hence defined as:

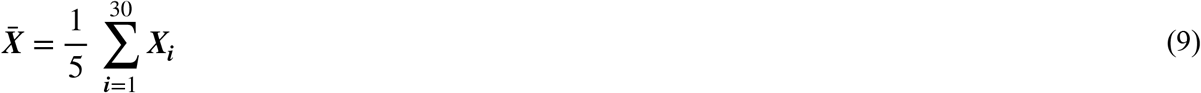

Where 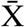 is the average value of the metrics after thirty executions and X_i_ is the metrics obtained after each execution.

The evaluated metrics are defined below:

#### Accuracy

It is calculated as the number of right predictions divided by the total number of predictions made across all classes. This is the proportion of correct classifications that a trained machine-learning model achieves. It is defined as:

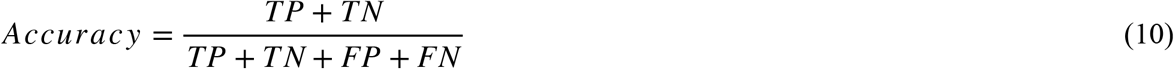

#### Precision

The ability of a classification model to locate just relevant information elements. Precision can be defined mathematically as the ratio of true positives to the sum of true positives and false positives. Here we have calculated the precision of predicting cancer samples correctly.

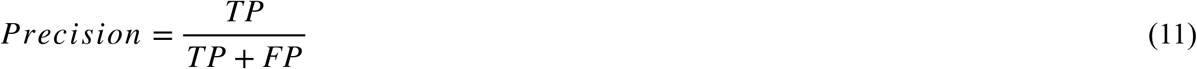

#### Sensitivity

Sensitivity quantifies the ability of a machine learning model to identify positive examples. Stated differently, it quantifies the probability of getting a positive outcome from a test. It is mathematically equal to the recall of the model.

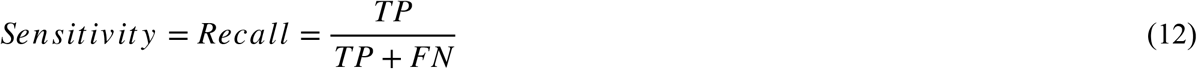

#### Specificity

The percentage of True negatives that the model accurately identifies is measured by specificity. It is also called the True Negative Rate (TNR). The model’s increased Specificity indicates that it correctly identified the majority of the negative outcomes. A low specificity value suggests that the model misinterpreted the negative outcomes as positive.

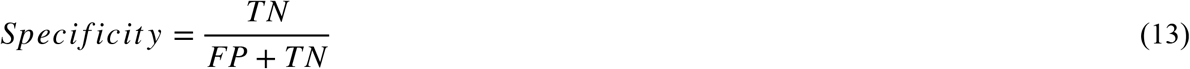

In all the above equations TP denotes True Positive, TN denotes True Negative, FP denotes False Positive and FN denotes False Negative.

#### ROC Curve

It is plotted with the True Positive Rate (TPR) on the y-axis and the False Positive Rate (FPR) on the x-axis. The ROC curve gives the tradeoff between True Positive Rate and a False Positive Rate at different threshold values. The ROC curve for an ideal classifier is a diagonal line from the bottom left corner to the top right corner. The area under the ROC curve is known as the Area Under the Curve (AUC). It provides an aggregate measure of performance across all possible classification thresholds. AUC is scale-invariant. It measures how well predictions are ranked, rather than their absolute values. It measures the quality of the model’s predictions irrespective of what classification threshold is chosen. The ROC curves were drawn to evaluate the performance of the models for five iterations.

## 4. Results

### 4.1 Feature Extraction Using Lasso Regression Model and Model Performance

The preprocessed dataset was split into training and validation set and the Lasso regression module was trained. The optimal value of was found to be 1.0 × 10^−6^ [Figure 3]. The confusion matrix of the model and the ROC curve were built using the sklearn Confusion Matrix package. The accuracy of the model was evaluated using the accuracy_score module of the sklearn library. The model was evaluated for 30 iterations with different training and testing sets. The model obtained an average accuracy of 84.58%, an average specificity of 89.75%, an average sensitivity of 83.28% and an average precision of 87.10% [Table 1]. We are not using this classifier as the primary classifier, this classifier model is used for ranking the features (genes) and setting the coefficient of insignificant genes to zero. The ROC curve was plotted along with the confusion matrix [Figure 4]. The average AUC value is 0.802 [Figure 4]. The absolute values of the coefficients of all the genes were extracted and the genes with 0 coefficients were omitted. After omission, the final gene count came down to 2527 from the initial figure of 16382. Hence the dimension of the final dataset containing the significant features is (51 2527) so a reduction of 13855 can be reached.

**Table 1:**
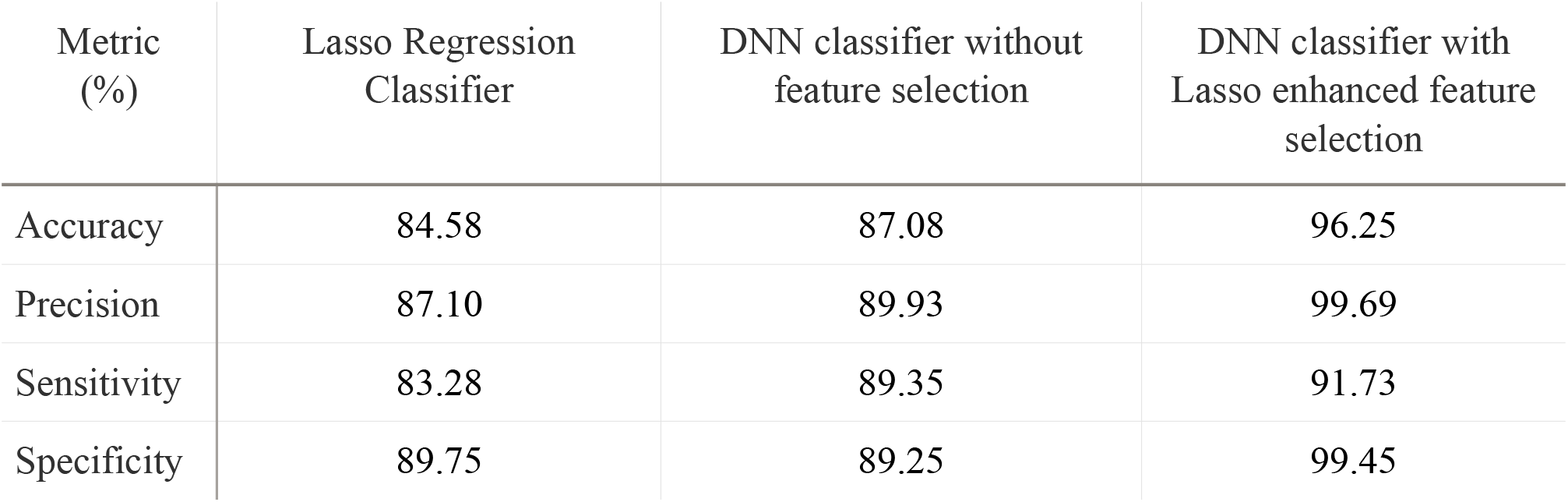
Validation metrics of the different models. All the models were evaluated for 30 iterations with different combinations of testing and validation sets and the average of the metrics were taken.

**Figure 3.**
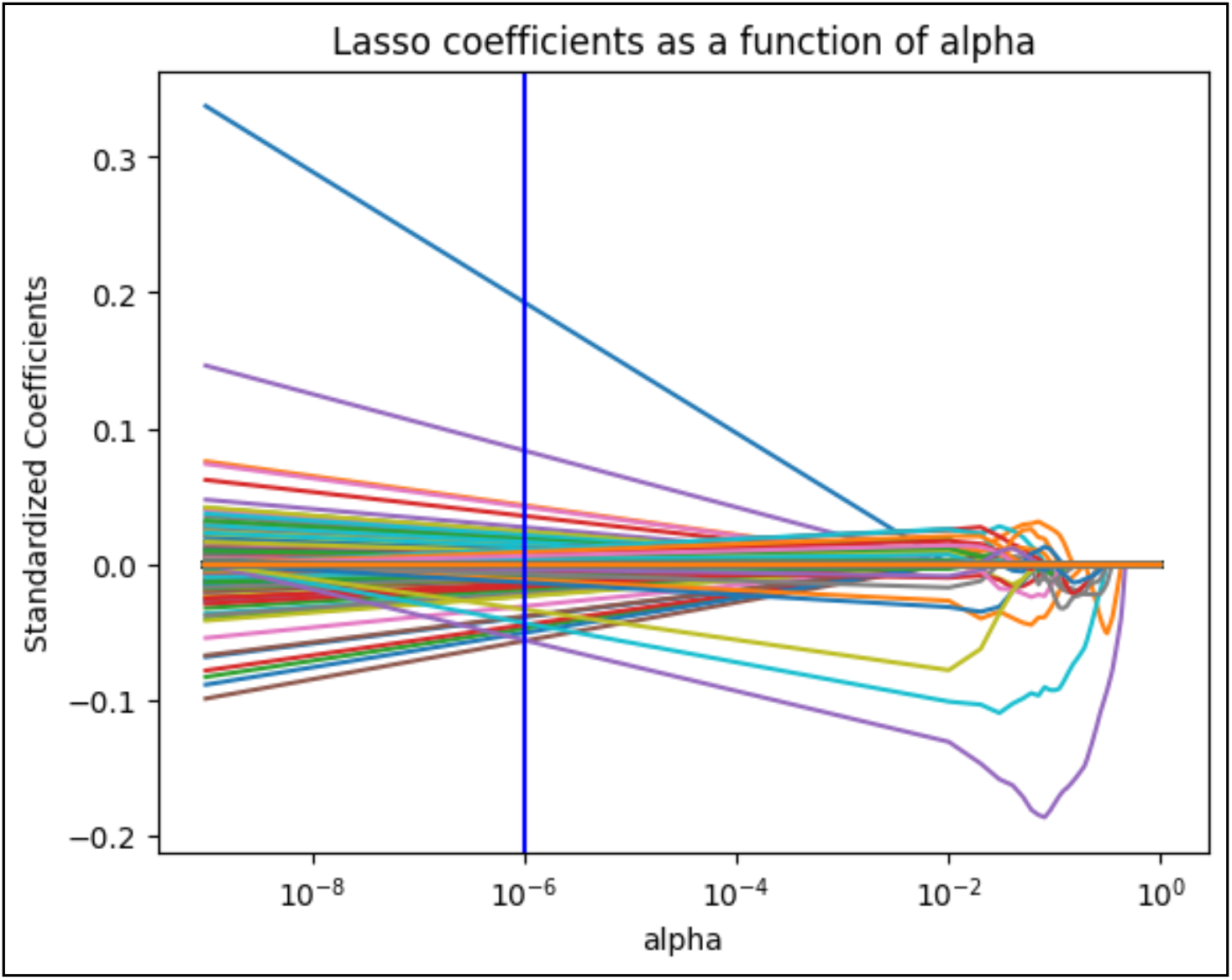
Plot of the Lasso cofficients of the different features(genes) as a function of alpha. The chosen alpha value of 1e-6 is shown with the vertical blue line.

**Figure 4.**
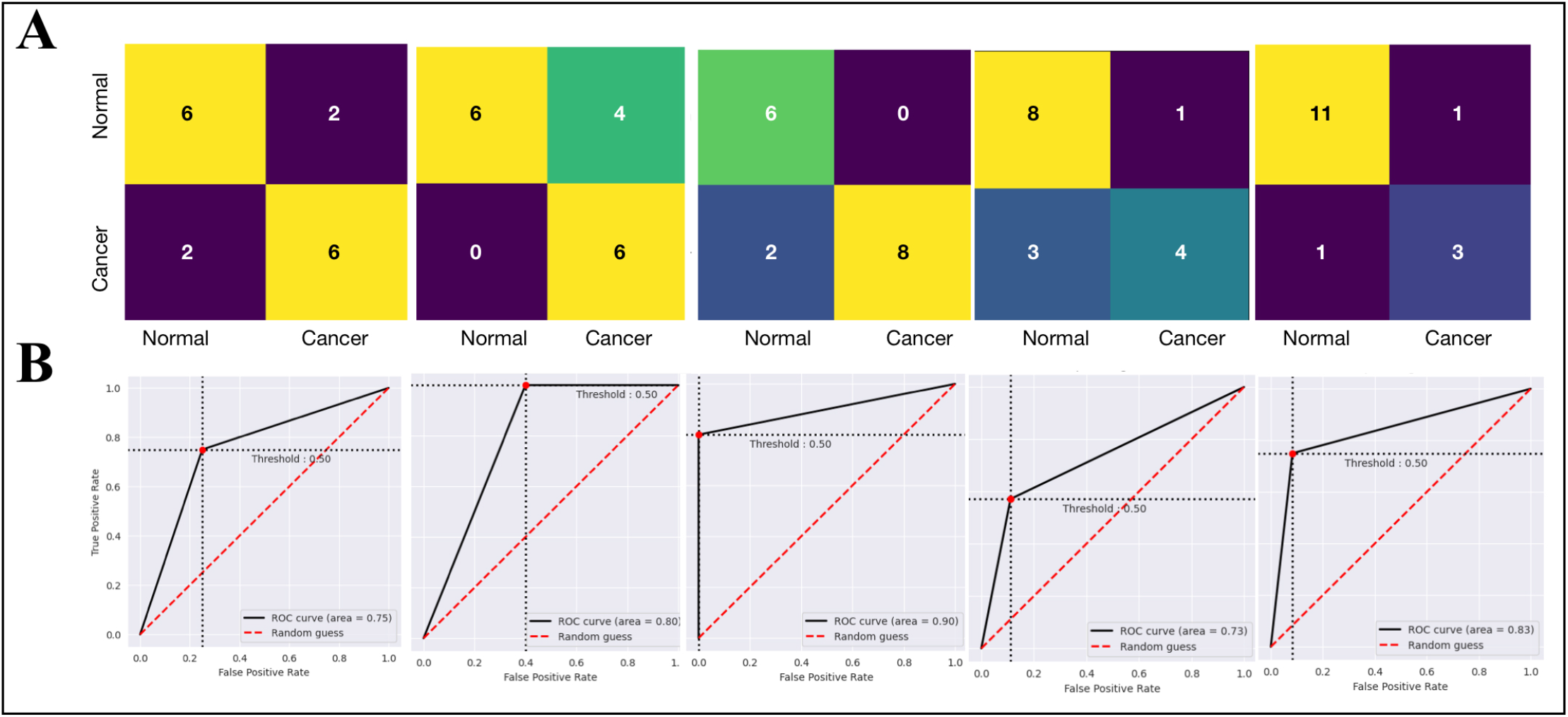
Model Evaluation of the Lasso regression classifier. (A) Confusion matrix representation of the first five iterations. (B) ROC curves corresponding to the first five iterations. The corresponding AUC scores are also shown.

### 4.2 Classification Using the Convergent Deep Neural Network for both primary and feature extracted datasets

In the first case, the primary dataset was split into training and validation sets and was fed into the DNN classifier which was trained [Figure 5] and its performance was evaluated for thirty iterations. The Confusion Matrix and the ROC curve for the first five iterations are shown in Figure 6. The model metrics were calculated for the thirty iterations. The average accuracy obtained was 87.08%, the average specificity was 89.25%, the average sensitivity was 89.35%, and the average precision was 89.93%. The average AUC score for the first five iterations was 0.876 [Figure 6(A)].

**Figure 5.**
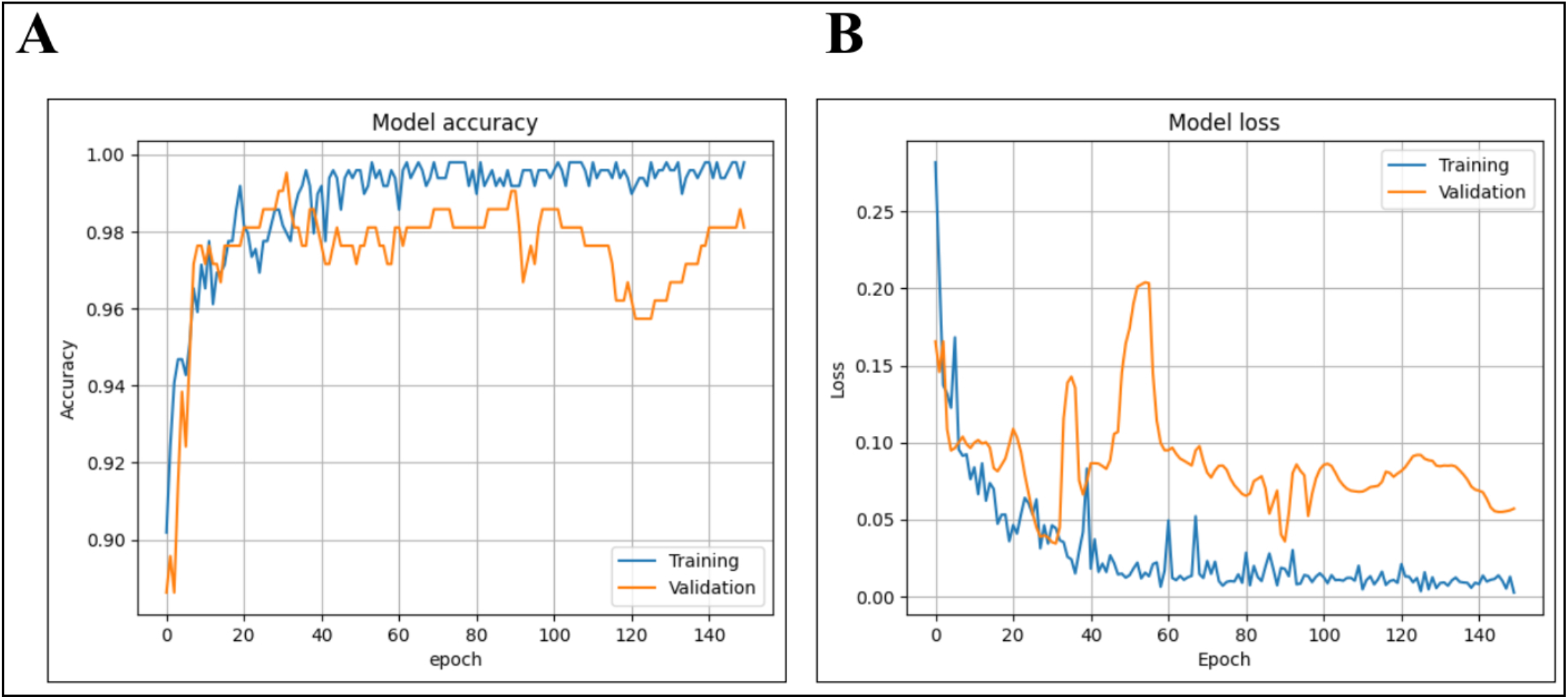
Accuracy versus number of epoch plot of the Deep Neural Network as obtained during training. (A) The accuracy curve corresponding to the training and validation. (B) The loss curve corresponding to the training and validation

**Figure 6.**
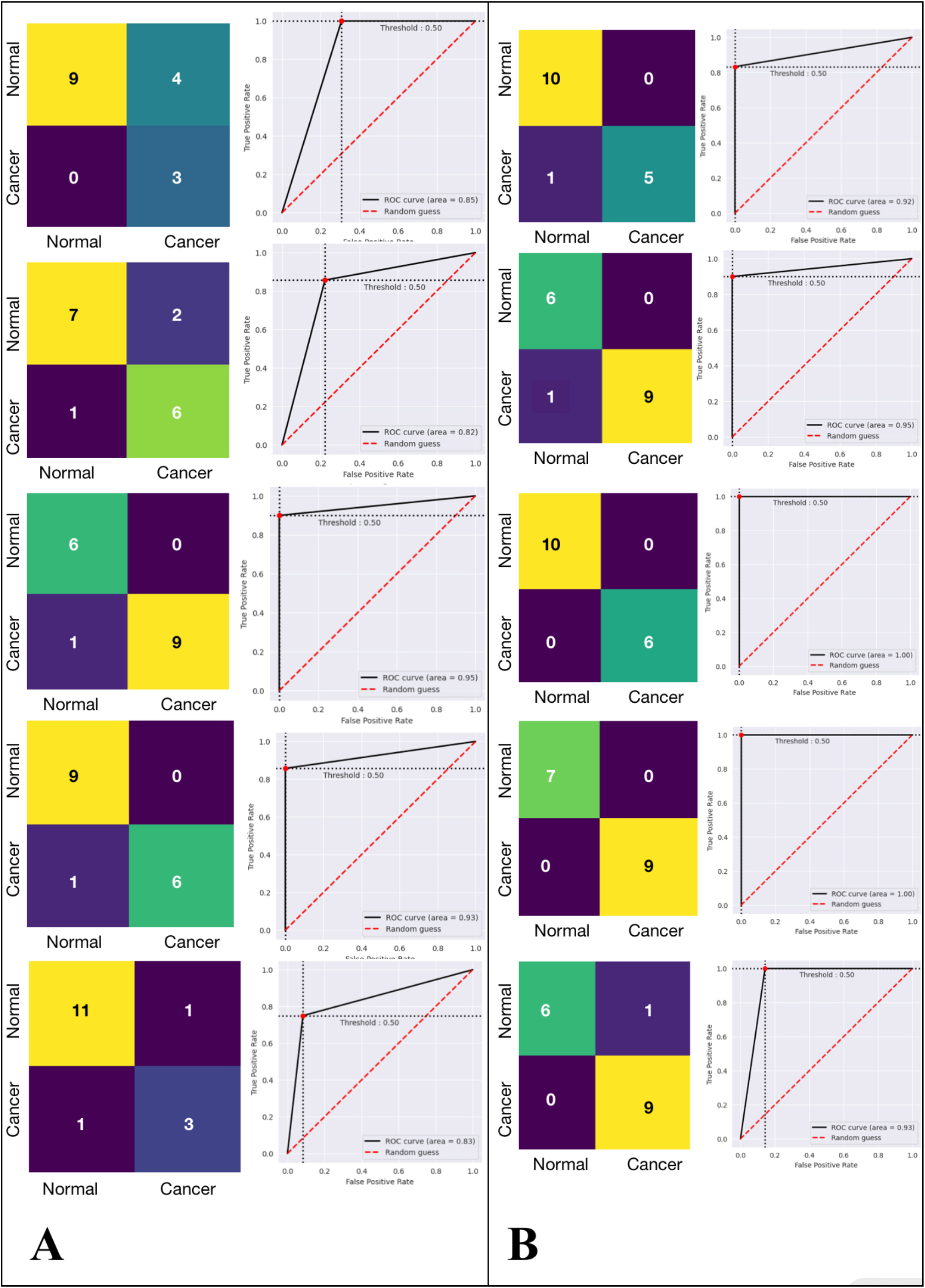
Model Evaluation of the Lasso regression classifier. (A) Confusion matrix and corresponding ROC curve representation for the first five iterations of the DNN classifier using the primary dataset. (B) Confusion matrix and corresponding ROC curve representation for the first five iterations of the DNN classifier using the feature selected dataset (proposed). The corresponding AUC scores are also shown.

Now in the second case, the feature-reduced dataset obtained in section 4.1 was taken. It was divided into training and validation sets. Like the first case, the performance of the DNN classifier was evaluated over thirty iterations. The average accuracy obtained was 96.25%, the average specificity was 99.45%, the average sensitivity was 91.73%, and the average precision was 99.68%. The average AUC score for the first five iterations was 0.96 [Figure 6(B)].

## 5. Discussion

We compared the results obtained between the three methodologies. The classification metrics are given in Table 1. We can see that with the initial dataset, the classification parameters show significant improvement with the use of the Convergent Deep Learning module as compared to the traditional Lasso Regression classifier. Now after feature selection and dimensionality reduction of the dataset and hence by combining the Lasso feature selection with the Convergent Deep Learning module, the classification parameters improved marginally with high significance[Fig 7]. We also compared the classification accuracy with some other classifiers designed in earlier works, the result of which is tabulated in Table 2.

**Table 2:** Comparison of the obtained result with earlier works. It should be noted that the accuracy of most of the models were not taken as. An average but the best was taken as the overall accuracy. In our case we took the average for thirty independent iterations. The proposed model can obtain accuracy of 100%.

| Reference | Classifier | Accuracy (%) |
| --- | --- | --- |
| [21] | SVM | 65.0 |
| [22] | SVM (linear) | 94.22 |
| [23] | Artificial Algae Algorithm enhanced SVM | 95.71 |
| [24] | CNN | 95.77 |
| Proposed model | Lasso enhanced CNN | 96.25 |

**Figure 7.**
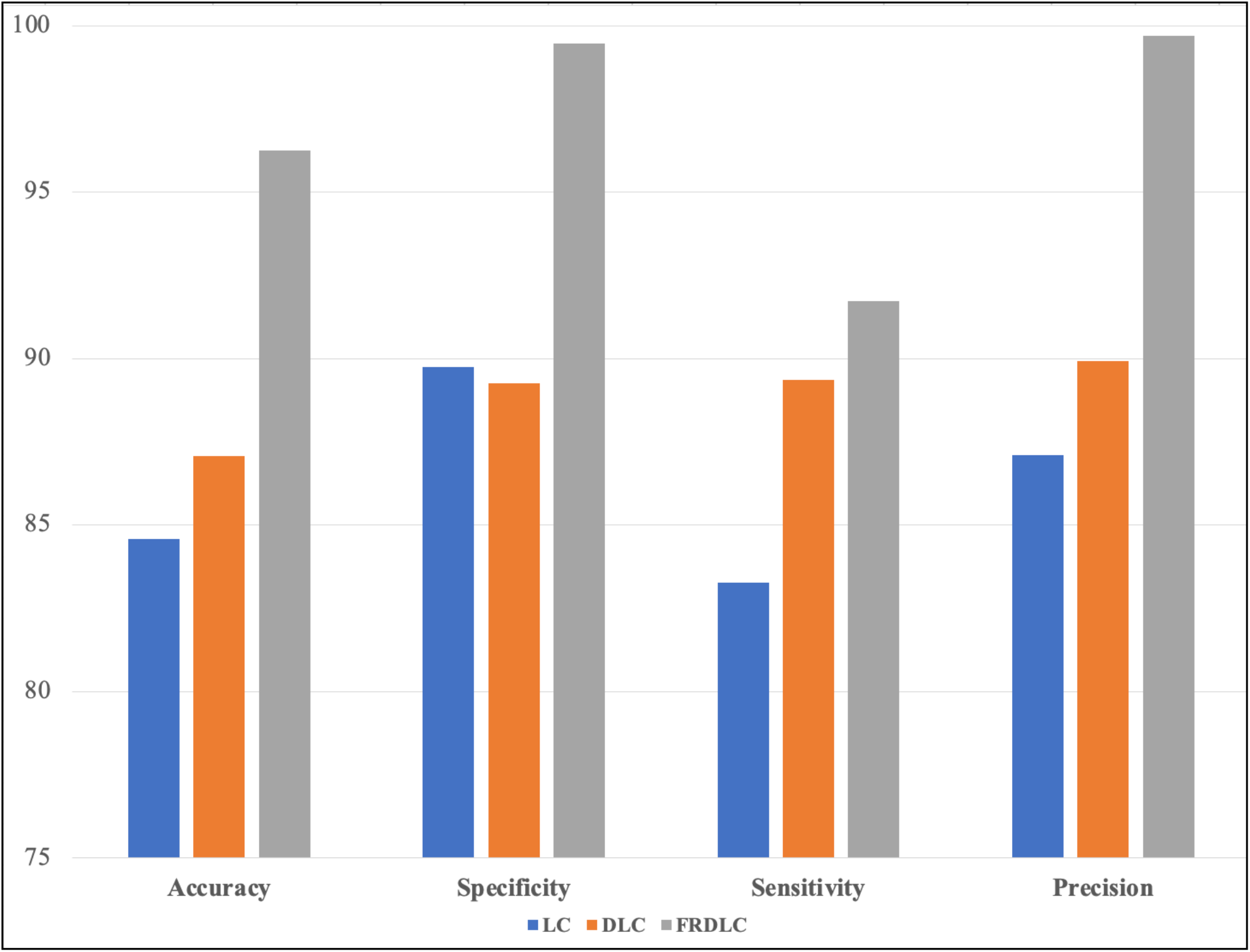
Performance comparison of the different classifier models. LC denotes Lasso Recression classifier, DLC denotes the Convergent Deep Learning Model without feature selection FRDLC denotes the Feature reduced Convergent Deep Learning Model (proposed).

From the result, it can be observed that feature selection can statistically improve the classification accuracy of the classifiers [Figure 8]. DNN follows highly efficient ML algorithms but a high dimensional dataset could result in noise and improper pattern recognition, which could limit its capabilities. Feature selection hence can be instrumental in reducing the number of input features and hence the dimensionality of the dataset. This could marginally improve the performance of the classifier.

**Figure 8.**
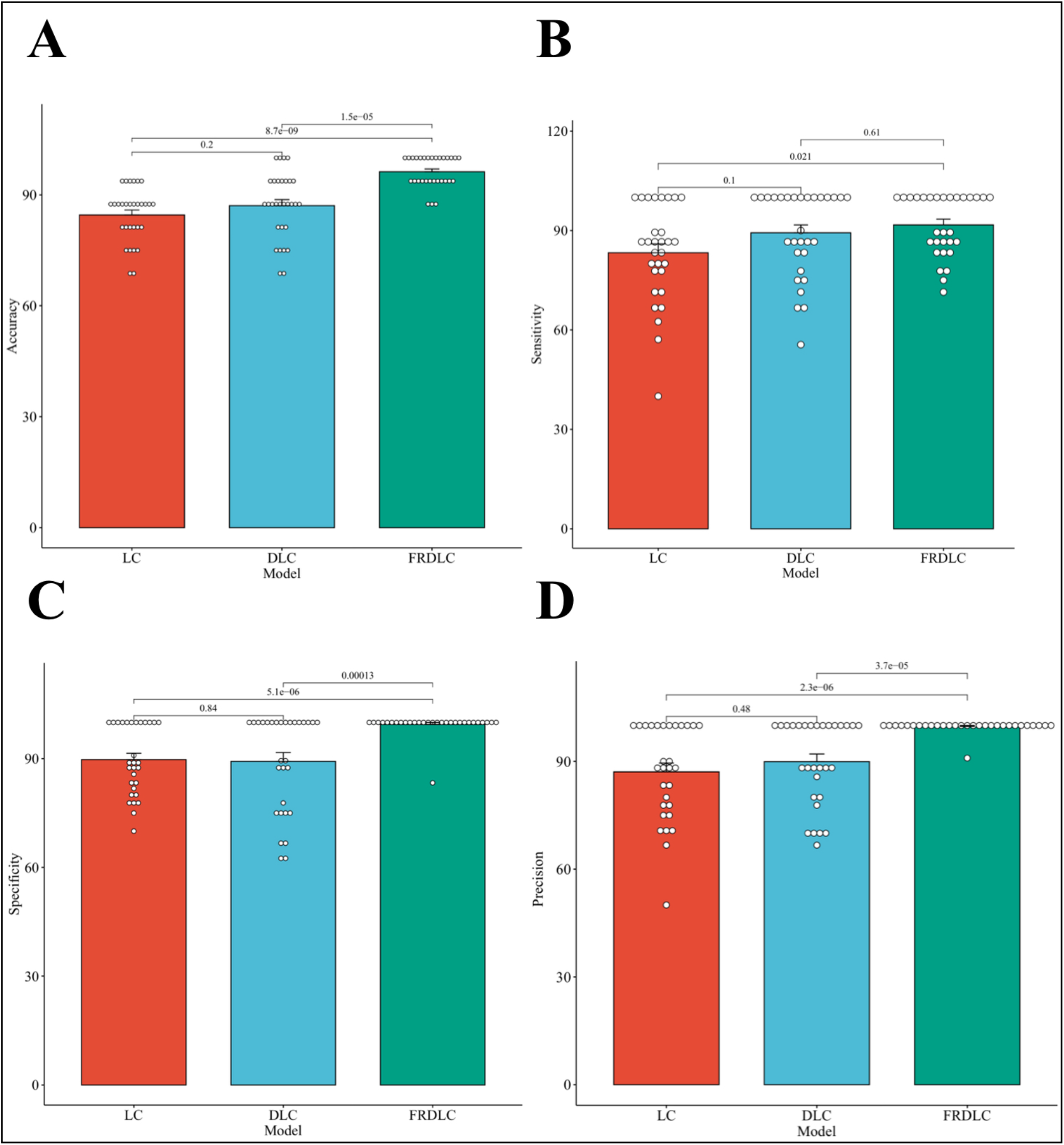
Significance comparison of the different classifier models by t-test. LC denotes Lasso Recression classifier, DLC denotes the Convergent Deep Learning Model without feature selection FRDLC denotes the Feature reduced Convergent Deep Learning Model (proposed). It is clearly demonstrated that Feature selection enhances the performance of the DNN classifier significantly.

## 6. Conclusion

Genetic diseases are one of the most significant health crisis all over the globe. These diseases are affecting a huge number of people. In most cases, the key to a better prognosis is early diagnosis. Techniques like microarray are hence gaining significance for probing these genetic abnormalities. This current work focuses on the classification of lung cancer samples using a hybrid DNN classifier enhanced with feature extraction capabilities of a Lasso Regression model. The model performance was improved by the addition of the Lasso feature extraction module. DNN classifies the data using its unique data-driven learning approach. While this approach works well, it is limited by the presence of a large amount of noise in the gene expression data, the repeatability of the results, and individual differences due to age, gender, genotype, stage of illness, and other variables. This disadvantage can be removed by implementing a statistical meta-analysis of combined different studies which would enable us to find the distinct disease signatures that are typically consistent in several studies [25]. The benefit of using Deep Learning is the improved classification accuracy combined with minimal computational time. Feature selection further improved the performance of the DNN classifier with the accuracy of prediction improving by 9.17%. Other than diagnosis this model can be also used for segregation between cancer types of any specific cancer which is also a significant problem. In the future improvements can be made by optimization of the network structure. This model can also detect effectively the presence of any gene mutations which could support research in the areas of genetics and virology. This model can be tuned for classification with any biological dataset.

## 7. Funding

None

## 8. Data Availability

The datasets used in this study is available in the NCBI Gene Expression Omnibus (GEO) with the accession ID GSE7670. All the codes used in the current work can be provided on request.

## 9. Declarations

### 9.1 Ethics approval and consent to participate

Not Applicable.

### 9.2 Consent for Publication

Not Applicable.

### 8.3 Competing interests

The author declares that he has no competing interests.

## Notes

### Competing Interest Statement

The authors have declared no competing interest.

